# Micro-engineered Konjac Glucomannan-Montmorillonite Hybrids as Multifunctional Biomaterials for Addressing Diet-Induced Obesity in Mice

**DOI:** 10.64898/2026.01.22.701163

**Authors:** Amin Ariaee, Hannah R. Wardill, Alex Hunter, Anthony Wignall, Amanda J. Page, Clive A. Prestidge, Paul Joyce

## Abstract

The growing prevalence of obesity necessitates innovative treatments. This study investigates a spray-dried konjac glucomannan–montmorillonite (KGM-MMT) hybrid designed to combine the fermentable, satiety-promoting effects of KGM with the lipid-binding and anti-inflammatory properties of MMT. In HFD-fed mice treated for 42 days with 2% w/w KGM-MMT, body weight gain was reduced by 7.6%, with an AUC of 5094[±[52.95, compared to 5513[±[81.35 in HFD controls (p < 0.0001). Serum IL-6 concentrations were reduced by 97% (p = 0.0002), while blood glucose decreased by 46% (p < 0.0001), outperforming reductions seen with MMT (24%, p = 0.0271) and KGM (16%, ns). Gut microbiota profiling demonstrated a significant 6.2-log[ fold increase in *Lactobacillaceae* (p = 0.023) and a 2.4-log[ fold increase in *Enterococcaceae* (p = 0.015) with KGM-MMT treatment. Predicted functional shifts revealed a 1.9-fold increase in short-chain fatty acid synthesis pathways and a 5.4-fold increase in bile acid deconjugation. Although the KGM-MMT hybrid did not consistently outperform its individual components in all measurements within the current study, it generally consolidated their metabolic benefits within a single dosage form. These findings support the utility of spray-dried KGM-MMT as a gut-targeted dietary strategy with additive effects on metabolic health. Future studies should explore underlying mechanisms and dosage effects of the hybrid formulation.

**Graphical abstract:** 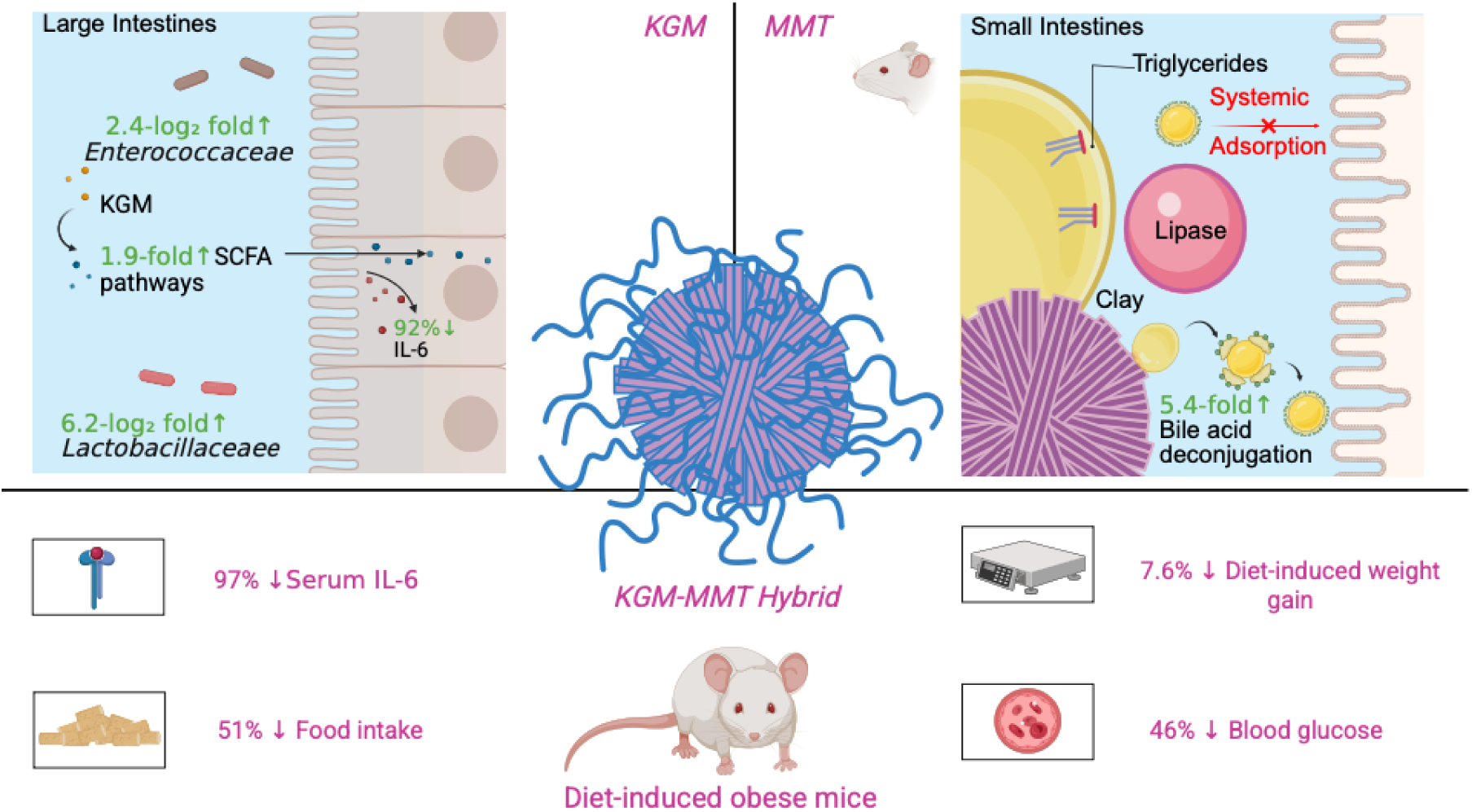

**Highlights:**

- Spray-dried KGM-MMT reduced HFD-induced weight gain by 7.6% in obese mice
- Serum IL-6 and glucose levels decreased by 97% and 46%, respectively
- 6.2-log[J and 2.4-log[J increases in *Lactobacillaceae* & *Enterococcaceae* relative abundance
- Bile acid deconjugation and SCFA pathways increased 5.4- and 1.9-fold
- KGM-MMT microparticles offer additive gut-targeted benefits in metabolic disease

## 1. Introduction

Obesity and its associated metabolic disorders continue to pose significant global health challenges, necessitating innovative therapeutic strategies to mitigate their impact. A growing body of evidence suggests that dietary interventions targeting the gut microbiota and systemic inflammation may hold promise in addressing these conditions ^1, 2^. Konjac glucomannan (KGM), a dietary fiber derived from the tubers of *Amorphophallus konjac*, has received 2 attention for its prebiotic and anti-inflammatory properties ^3–5^. Its high viscosity and gel-forming properties make it effective for weight management and reducing nutrient absorption, which contribute to its anti-obesity effects ^6^. Additionally, KGM has been demonstrated to selectively enhance the gut microbial taxa that are associated with improved gut barrier function, protecting against *metabolic endotoxemia* ^1, 7^.

Beyond its dietary uses, KGM has been incorporated into biomaterials, such as hydrogels and wound dressings, leveraging its biocompatibility and structural properties to support tissue regeneration and drug delivery applications ^8^. However, despite its widespread use in dietary and biomedical applications, KGM’s potential in addressing chronic systemic inflammation, a hallmark of obesity and metabolic diseases, remains underexplored ^9^. Further, the high viscosity of unmodified KGM powder lends itself to choking hazards due to its capacity to swell up to 50 times its weight in aqueous conditions, such as the stomach, posing significant dosing challenges ^10^. Hybridizing KGM with complementary materials into micron-sized particles offers a promising strategy to enhance its therapeutic efficacy. This approach reduces particle size and improves uniformity, mitigating risks associated with traditional KGM dosing while preserving and combine its therapeutic effects with other biomaterials.

Adsorbent clays, such as montmorillonite (MMT), provide promising avenues for such applications. MMT, a naturally occurring layered silicate, is known for its excellent adsorption properties, biocompatibility, and ability to modulate the gut microbiota and alleviate inflammation during diet-induced obesity ^11–13^. Its unique structural properties allow MMT to self-assemble into porous 3D microstructures, which are considered optimal for maximizing its adsorption capacity ^13, 14^. This porous network enhances its capacity to bind and remove toxins, and these adsorbent properties have more recently been investigated for the removal of lipolytic products from the intestinal lumen, reducing the metabolic effects of diets high in fats ^12, 13, 15, 16^. A previous study demonstrated that spray dried MMT adsorbed 42% of all lipid species in an *in vitro* lipid digestion model, whilst reducing diet-induced weight gain in rats similarly to orlistat, a pancreatic lipase inhibitor used as an anti-obesity drug ^13^.

As such, the current study investigates the additive effects of micro-structured KGM-MMT hybrids. This innovative approach aims to combine KGM’s prebiotic and anti-inflammatory properties with MMT’s adsorption capabilities through a hybrid material manufactured using the process of spray drying. By leveraging MMT’s porous 3D microstructures and KGM’s fermentable polysaccharide profile, the hybrid formulation is designed to target both the restriction of dietary fat digestion and promotion of the gut microbiota, addressing key drivers of metabolic dysfunction. This is the first study to examine spray-dried KGM-MMT hybrids that concurrently restrict diet-induced weight gain and gut microbiota modulation as a multi-mechanistic oral intervention for metabolic diseases.

## 2. Methods and Materials

### 2.1. Materials

KGM was purchased from Sigma-Aldrich (Castle-Hill, Australia). MMT was supplied by Pharmako Biotechnologies (Frenchs Forrest, Australia). Medium chain triglycerides (MCT, Miglyol 812) with caprylic acid and capric acid at 50/50 %wt , were purchased from Hamilton Laboratories (Adelaide, Australia). 6 to 8-week-old C57BL/6 mice were acquired from the breeding program at the South Australian Health and Medical Research Institute Centre (Adelaide, Australia). All chemicals and solvents used in this study were of analytical grade and ultra-pure water (Milli-Q) was utilized throughout the experimental procedures.

### 2.2. Fabrication of KGM-MMT hybrids via spray drying

KGM-MMT hybrids were prepared using a spray drying technique, following previously established methods ^17^. Briefly, an aqueous dispersion of KGM at 2% w/v was formed by dispersing 20 g of KGM in 1 L of Milli-Q water. The solution was stirred at room temperature for 30 minutes. Simultaneously, 20 g of MMT was dispersed in 1 L of Milli-Q water under similar conditions. The KGM and MMT dispersions were then combined in a 1:1 mass ratio.

The resulting mixture was spray dried using a mini Spray-Dryer B-290 (Büchi, Switzerland) under the following conditions: inlet temperature of 200°C, outlet temperature of 105°C, aspirator setting at 100%, nozzle cleaning setting at 9, compressed air flow rate of 40 mm, and product flow rate of 7 mL/min.

### 2.3. Morphological analyzing via Scanning Electron Microscopy (SEM)

The size and morphological characteristics of the fabricated KGM and its precursor materials were analyzed using a Zeiss Crossbeam 540 scanning electron microscope (Oberkochen, Germany). For SEM sample preparation, a small quantity of each sample was adhered to carbon tape mounted on SEM stubs. The samples were then coated with a platinum layer approximately 10–20 nm thick using a sputter coater. Imaging was performed at an accelerating voltage of 1–2 kV. SEM micrographs were processed and analyzed with AztecOne software (Oxford Instruments, Abingdon, United Kingdom).

### 2.4. Particle size analysis

The particle size of KGM, MMT, and spray-dried KGM-MMT hybrids was determined using laser diffraction (Mastersizer, Malvern Instruments, UK). Samples were dispersed in an aqueous buffer at pH 6.5, simulating the pH conditions of the small intestines, under continuous stirring for 60 min. The particle size distribution was characterized by the D_50_ value, which represents the median particle diameter where 50% of the particles by volume are smaller than this size. Measurements were performed in triplicate.

### 2.5. In vitro simulated gastrointestinal lipolysis assay

An in vitro intestinal lipolysis model was used to evaluate the effect of KGM, MMT, and spray-dried KGM-MMT hybrids on lipid digestion under simulated fasted and fed state simulated intestinal conditions (pH 6.5) ^13, 14^. Briefly, the simulated fasted and fedstate gastric (pH 1.6) and intestinal media (pH 6.5) were prepared using a 50 mM Tris–maleate buffer. Media were freshly prepared and used within 48 hours. Pancreatin extract was prepared by dissolving 2 g of pancreatin powder in 10 mL of the respective media (Fasted state simulated intestinal fluid (FaSSIF) or fed-SSIF (FeSSIF), pH 6.5) and centrifuging at 2268 × g for 20 minutes at 4 °C. The supernatant was collected and stored at 4 °C until use. The digesting lipid, MCT (625 mg) was dispersed in 20 mL of FaSSIF or FeSSIF by continuous stirring at 600 rpm for 10 minutes in a thermostated glass reaction vessel maintained at 37 °C. KGM, MMT, or spray-dried KGM-MMT hybrids were incorporated into the dispersion at 10% w/w relative to the lipid content. The pH of the lipolysis media was adjusted to 6.5 ± 0.01 using 0.1 M sodium hydroxide or hydrochloric acid. Lipolysis was initiated by adding 2 mL of pancreatin extract (equivalent to 2000 tributyrin units) to the reaction vessel. The liberation of free fatty acids (FFAs) was monitored for 25 minutes using a pH-stat titrator (902 Titrando, Metrohm, Switzerland), which maintained a constant pH of 6.5 by titrating 0.6 M sodium hydroxide.

### 2.6. *In vivo* study design

The *in vivo* study was approved by the Animal Ethics Committee at the South Australian Health and Medical Research Institute Centre and adhered to the NIH Principles of Laboratory Animal Care (NIH publication #85-23, revised 1985), the Australian Code for the Care and Use of Animals for Scientific Purposes (8th edition, 2013, revised 2021), and ARRIVE 2.0 guidelines for *in vivo* research ^18^. A well-established high-fat diet (HFD) model was used, involving 40 male C57BL/6 mice randomly assigned to four groups (n = 10 per group) via computer-generated allocation to minimize selection bias. The groups included a control HFD group (44% kcal from fat) and three treatment groups, where 2% w/w KGM, 2% w/w spray-dried MMT, or 2% w/w spray-dried KGM-MMT hybrids were incorporated directly into the HFD, which the mice had free access to throughout the 6-week study. Mice were housed in groups of three under standardized conditions (12-hour light/dark cycle, 22 ± 2°C, 55 ± 10% humidity) for a one-week acclimation period before a 42-day treatment phase. In the final week of the study, mice were individually housed in metabolic monitoring cages (Promethion, Sable Systems International, United States) to precisely measure variables such as respiratory quotients, energy expenditure, and food intake during a 24h period for 3 days at the end of the treatment phase. These specialized cages were equipped with high-resolution sensors for continuous monitoring of metabolic activity and energy expenditure over the final 24h period before the end of the treatment phase. Food intake patterns were tracked with automated feeding systems that recorded meal sizes and feeding frequency, while integrated motion sensors captured movement and activity levels. The cages were designed to minimize environmental stress and ensure consistent conditions, including controlled temperature, humidity, and airflow.

Body composition was measured using EchoMRI to obtain accurate and non-invasive assessments of fat mass, lean mass, and total body water. At the end of the treatment phase, mice were anesthetized with isoflurane. Approximately 1 mL of blood was collected by cardiac puncture for serum analysis, and mice were then euthanized by cervical dislocation.

### 2.7. 16s rRNA gene sequencing of fecal samples

Following humane euthanasia, fecal samples were collected and shipped to the Australian Genomics Research Facility (Brisbane, Australia) for DNA extraction and 16S ribosomal RNA (rRNA) gene sequencing targeting the V3–V4 hypervariable regions. The resultant sequences were clustered into operational taxonomic units (OTUs) at a 97% similarity threshold using QIIME 2 and the Silva reference database (Release 138.1) [29, 30]. OTU taxonomic assignments were determined with QIAGEN CLC Genomics Workbench Version 23.0.4 and the QMI-PTDB TaxPro index (June 2021) [31]. Microbial diversity was characterized by calculating Shannon, Chao1 indices and total OTU number for alpha diversity. Beta diversity indices were obtained using Bray-Curtis dissimilarity, Jaccard indices, and weighted/unweighted UniFrac distances. Statistical differences in beta diversity were evaluated via Permutational Multivariate ANOVA (PERMANOVA), and PCoA plots for each group were displayed with 95% confidence ellipses to visualize group clustering and variability [34]. The exclusion of mice samples was due to them not meeting data quality control parameters as set by the Qiagen Metagenomic module. This resulted in a final sample size of n=8 for each group in the study.

### 2.8. 16s rRNA metagenomic predictions using PICRUSt2

Phylogenetic Investigation of Communities by Reconstruction of Unobserved States (PICRUSt) version 2.0 was integrated as a QIIME2 plugin to predict the presence and abundance of enzymes and metabolic pathways within each sample. Enzyme predictions were generated using the Enzyme Commission (EC) database (Nov-2023), and the corresponding EC numbers were subsequently applied to estimate metabolic pathway abundances via the MetaCyc Pathway Database (May-2022).

### 2.9. Serum proinflammatory cytokine quantification using ELISA

Following centrifuging whole blood at 2000 g for 20 minutes at 4°C, the serum fraction was carefully collected and stored at –80°C until further analysis. Serum interleukin-6 (IL-6) and Interferon-gamma (IFN-γ) concentrations were measured using ELISA kits (ThermoFisher Scientific, Australia), following the manufacturer’s instructions. Quantification was carried out by generating standard curves from known IL-6 concentrations. All assays were performed in duplicates according to standardized protocols to ensure consistency and reproducibility. The limit of detection is reported as 3 pg/mL.

### 2.10. Statistical analysis

All experimental data, except for 16S rRNA gene sequencing results, were analyzed using GraphPad Prism Version 10.2.0 (Boston, MA, USA). Prior to statistical analysis, the normality of data distribution and residuals was assessed using the Shapiro-Wilk test. Where data deviated from normality, non-parametric Kruskal-Wallis tests with Dunn’s post hoc were employed. Where normality is found, statistical comparisons were made via a one-way ANOVA with Tukey’s post hoc for multiple comparisons. All data are presented as mean ± SD unless otherwise stated. Line and bar graph data are presented as mean ± standard deviation (SD), while box and whisker plots display the full data range (min-max). Statistical comparisons were conducted using one-way ANOVA with Tukey’s post hoc (or Kruskal-Wallis with Dunn’s post hoc when normality was not met). Asterisks represent significance between groups, where: * = p ≤ 0.05, ** = p ≤ 0.01, *** = p ≤ 0.001, and **** = p ≤ 0.0001.

## 3. Results and Discussion

### 3.1. Spray drying of KGM with MMT produces microparticles

The structural and morphological properties of KGM and MMT are critical to their functionality and therapeutic potential when combined as hybrids. The SEM analysis reveals key microstructures of KGM, MMT, and their hybrids, highlightling properties that may contribute to their enhanced performance. The SEM image of KGM powder shows amorphous particles with a porous morphology, a structure that allows it to form a network that can trap water effectively ^19^ (**Fig. 1A**). KGM’s high water-holding capacity enhances its solubility and viscosity, facilitating fermentation by gut microbiota. These properties contribute to its role as a fermentable dietary fiber, supporting microbial metabolism in the gut. As a result, KGM supplementation can proliferate the growth of commensal taxa associated with health benefits, whilst enhancing intestinal microbiota metabolites in short chain fatty acids (SCFA) ^20^. However, KGM’s particle size spans between 20-50 µm, which may limit its dispersion as well as its interaction with the microbiota in aqueous environments, such as the large intestines. Additionally, the high viscosity and swelling capacity of unmodified KGM powder pose risks of choking or esophageal obstruction, particularly when orally consumed in large doses ^21^. Similarly, an *in vitro* fermentation study demonstrated that reducing KGM’s particle size via acid modification resulted in significantly higher substrate utilization by the inoculated fecal microbiota and enhanced SCFA production ^22^. These findings highlight the importance of reducing KGM’s particle size to improve its safety and functionality. In contrast, MMT particles are irregular and compact, characterized by a rough surface morphology that reflects the extrusion of the layered platelet structure within the aluminosilicate clay (**Fig. 1B**).

**Figure 1.**
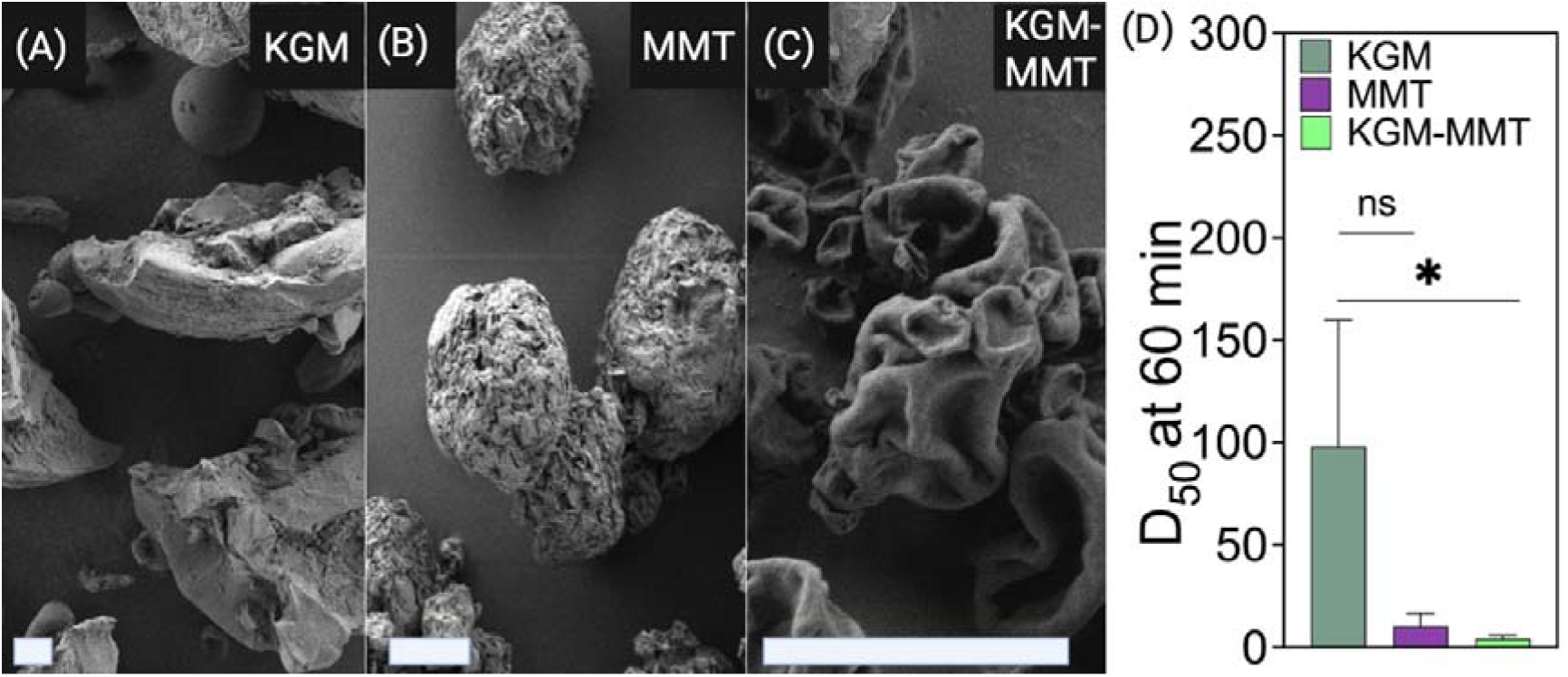
SEM images illustrate the morphological differences between KGM-MMT hybrids and their precursor materials. **(A)** KGM exhibits irregular, 20–50 µm-sized porous structures. **(B)** Spray-dried MMT forms 8–10 µm-sized spherical particles with a smoother surface, indicative of enhanced uniformity and dispersion. **(C)** Hybrid particles formed by spray drying KGM with MMT demonstrate KGM incorporation into the MMT matrix, resulting in 2–5 µm particles with sheet-like morphology. **(D)** Particle size analysis of spray-dried KGM-MMT dispersed in an aqueous pH 6.5 buffer reveals significantly smaller particle sizes compared to its individual precursors, highlighting the effect of spray drying on particle size reduction. Scale bars represent 10 µm. Particle size data is presented as mean ± SD (n = 3 per group). Statistical differences were determined using one-way ANOVA with Tukey’s comparisons post hoc (ns > 0.05; *p ≤ 0.05, **p ≤ 0.01, ***p ≤ 0.001, ****p ≤ 0.0001).

The spray-dried KGM-MMT hybrid exhibits a unique morphology with wrinkled, sheet-like structures, indicating a structural transformation induced by the spray-drying process (**Fig. 1C**). This transformation reduces particle size to 2–5 µm and enhances uniformity ^13, 14^. Particle size analysis further corroborates these findings, as shown in **Fig. 1D**. After 60 minutes of dispersion in an aqueous intestinal pH 6.5 buffer, the spray-dried KGM-MMT hybrid exhibited a median particle size of 4.19 ± 0.75 µm, corresponding to a 23-fold reduction compared to KGM (97.99 ± 30.00 µm) and a 2.5-fold reduction compared to unmodified MMT (10.29 ± 3.19 µm). Tukey’s multiple comparisons test confirmed that the difference between KGM-MMT and KGM was statistically significant (p = 0.0423), while the difference between KGM-MMT and MMT was not significant (p = 0.9764). These reductions highlight the effectiveness of spray drying in producing finer particles, corroborating with previous findings ^13, 16^. The smaller, more uniform particles increase the surface area, enhancing KGM-MMT’s gut activity. For instance, spray-dried smectite clays have been shown to reduce *in vitro* fat digestion by up to 1.4-fold compared to their unmodified forms ^13^. MMT is a well-documented adsorbent for lipolytic products, such as FFAs in the small intestine, due to its high cation exchange capacity and layered structure, which predominately bind FFAs via the silanol groups at the clay’s broken platelet edges ^13–16^. Spray drying optimizes these properties by improving dispersion in the gastrointestinal tract, enabling MMT to adsorb more lipolytic products and remove them from the small intestine before systemic absorption ^13^. The hybridization of KGM-MMT is postulated to offer a multi-faceted effect in the gastrointestinal tract: MMT adsorbs lipolytic products in the small intestine, while KGM reaches the large intestine intact, serving as a fermentable substrate for the residing microbiota. Previous studies have demonstrated that spray drying effectively hybridizes fermentable polysaccharide-based gut-actives, such as inulin-MCT, significantly reducing weight gain in obese rats ^23^. These findings establish a proof-of-concept for using spray drying to develop multifunctional biomaterials with enhanced therapeutic potential. Future studies should also assess the particle size behavior of such materials in pH buffers relevant to the gastric phase, in addition to the intestinal phase, to better understand their stability and functionality across different regions of the gastrointestinal tract.

### 3.2. KGM-MMT hybrids suppress in vitro simulated intestinal fat digestion

Under fasted conditions, the KGM-MMT hybrid significantly suppressed intestinal lipid digestion compared to KGM, MMT, and HFD controls (**Fig. 2A**). After 60 minutes of in vitro MCT digestion, the free fatty acid (FFA) release for KGM-MMT was 525.1 ± 2.7 µmol, representing a 4.0-fold reduction compared to HFD controls (1451 ± 1.2 µmol, p < 0.0001), a 2.71-fold greater reduction compared to KGM (1356 ± 2.1 µmol, p < 0.0001), and a 2.21-fold greater reduction compared to MMT (1211 ± 2.7 µmol, p < 0.0001). Dunn’s multiple comparisons test confirmed significant differences between all groups, with KGM-MMT showing the most profound inhibition (p < 0.0001 for all comparisons). The substantial decrease in FFA release achieved by KGM-MMT likely results from the additive effects of MMT’s lipid-adsorptive properties and KGM’s viscosity-enhancing capability, both of which reduce lipid accessibility to digestive enzymes ^16, 24^. Moreover, KGM particles may disrupt the lipid-in-water interface, impairing emulsification and thereby diminishing the surface area available for enzymatic action ^25^.

**Figure 2.**
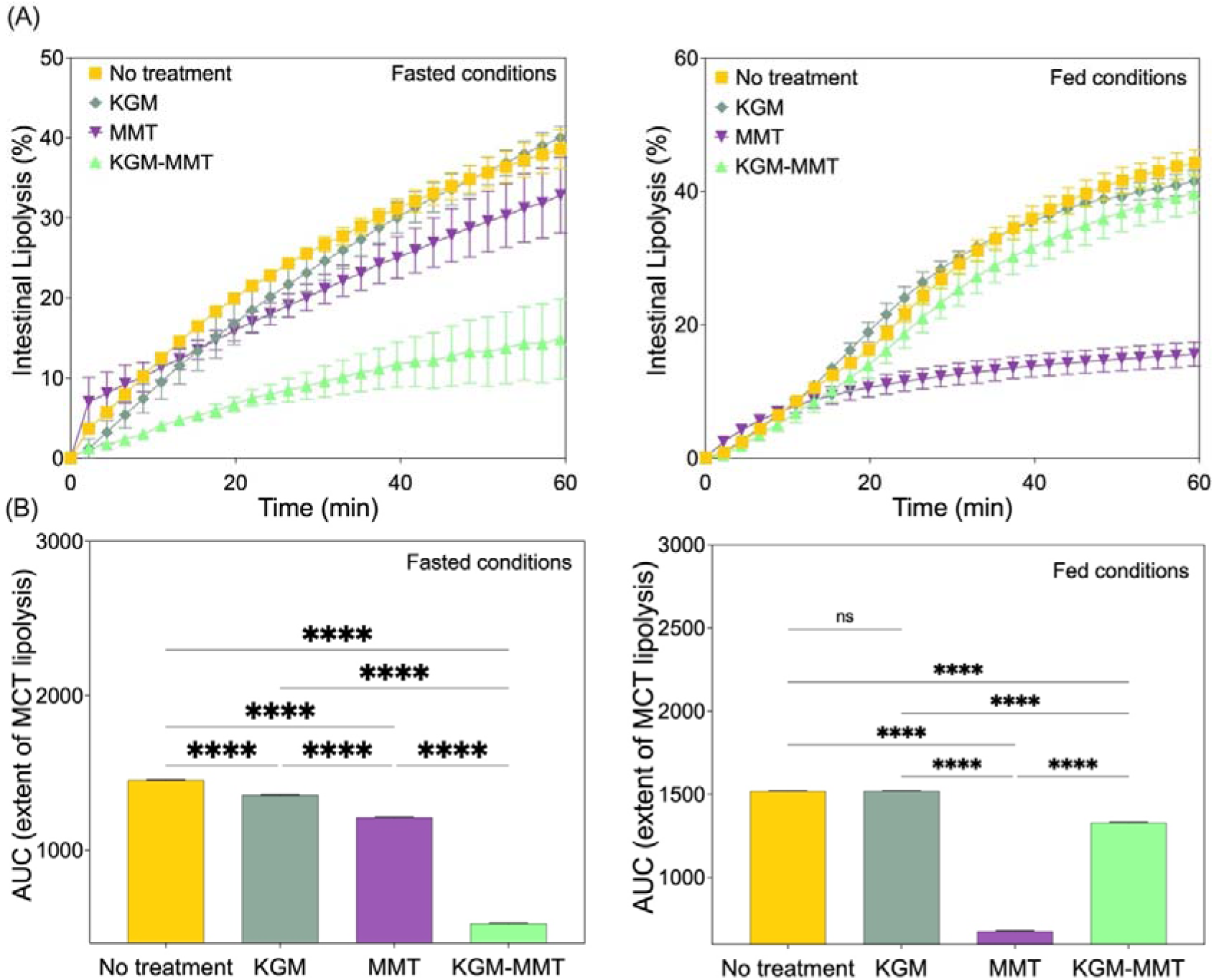
**(A)** *In vitro* intestinal lipolysis of MCT with KGM, MMT, and KGM-MMT hybrids under fasted (left) and fed (right) conditions. KGM-MMT hybrids show the greatest suppression of FFA release in the fasted state, while MMT is most effective in the fed state. **(B)** AUC analysis reveals reductions in lipolysis by KGM-MMT hybrids and MMT, with KGM having no effect in the fed state. AUC data is presented as mean ± SD (n = 3 per group). Statistical differences were determined using the Kruskal–Wallis test with Dunn’s multiple comparisons post hoc (ns > 0.05; *p ≤ 0.05, **p ≤ 0.01, ***p ≤ 0.001, ****p ≤ 0.0001)

In the fed state, a different pattern was observed. MMT alone was the most effective in reducing FFA release (678.3 ± 1.5 µmol), achieving a 1.9-fold reduction compared to HFD (1518 ± 1.5 µmol, p < 0.0001). In comparison, the KGM-MMT hybrid reduced FFA release by 1.14-fold (1329 ± 2.3 µmol, p < 0.0001), while KGM alone (1519 ± 1.0 µmol) had no significant effect (p = 0.8720). This suggests that under emulsifier-rich fed conditions, MMT’s lipid adsorption dominates due to increased micelle interactions ^5^, whereas KGM’s viscosity-inducing and emulsification-disrupting actions become less effective. The cumulative extent of lipolysis, as reported by the area under the curve (AUC), was reduced by KGM, MMT, and KGM-MMT by 6.5%, 17%, and 64%, respectively, in the fasted state (**Fig. 2B**). The KGM-MMT hybrid combines these mechanisms, resulting in a dramatic reduction in FFA release. Under fed conditions, MMT and KGM-MMT hybrids reduced FFA release by 55% and 12%, respectively, while KGM showed no effect. The dominance of MMT under fed conditions reflects its enhanced adsorption of lipolytic products in emulsifier-rich environments, while the hybrid’s reduced efficacy compared to the fasted state may result from its lower MMT content.

### 3.3. Restoration of the microbiota by KGM in HFD-fed mice

Significant changes in gut microbiota composition and diversity were observed following the inclusion of KGM-MMT hybrids and their precursors in the diet of HFD-fed mice over a 42-day treatment period. Dietary supplementation with KGM significantly increased alpha diversity indices, including Chao1 richness, Shannon’s index, and total observed species counts (**Fig. 3A–D**). KGM treatment improved these metrics by approximately 1.2- to 2.4-fold relative to the HFD group, with statistically significant differences observed for Shannon’s index (p = 0.0054), Chao1 richness (p = 0.0143), and total observed species (p = 0.0069). This enhancement in alpha diversity is crucial, as reduced gut microbial diversity is a recognized hallmark of gut dysbiosis and is strongly associated with metabolic disorders such as obesity, diabetes, and systemic inflammation ^26^. Lower alpha diversity, particularly reductions in Chao1 and Simpson indices, has been directly linked to poorer metabolic health profiles in obese individuals and children, indicating a compromised gut ecosystem that predisposes to metabolic diseases ^27^. Restoration of diversity through dietary fibers like KGM may therefore promote a more resilient and metabolically protective microbiome structure.

**Figure 3.**
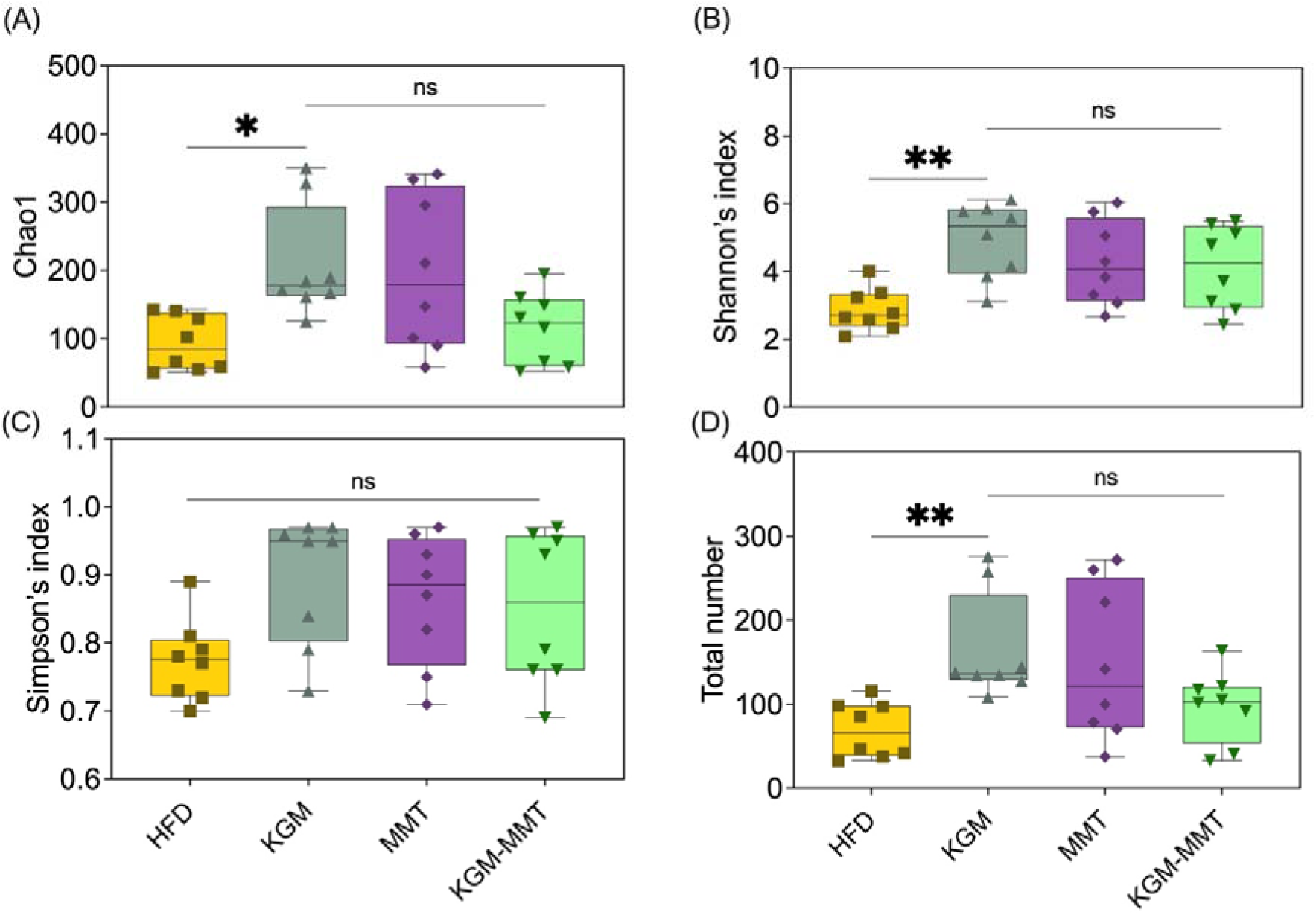
16S rRNA sequencing results highlighting microbial diversity and taxonomic composition across treatment groups. Alpha diversity indices, being (**A**) Chao1, (**B**) Shannon, (**C**) Simpson indices and (**D**) total number of operational taxanomic units, are presented as box and violin plots displaying minimum to maximum values (n = 8 per group), demonstrating the significant enrichment of microbial richness with KGM inclusion in the HFD. Statistical differences were determined using the Kruskal–Wallis test with Dunn’s multiple comparisons post hoc (ns > 0.05; *p ≤ 0.05, **p ≤ 0.01, ***p ≤ 0.001.

Increased microbial diversity supports greater ecosystem stability and resistance to pathogenic colonization, mitigating adverse outcomes such as increased intestinal permeability and systemic endotoxemia, which are triggered by Western-style, high-fat diets ^28^. Elevated levels of lipopolysaccharides (LPS) and decreased microbiota diversity contribute additively to systemic inflammation and metabolic dysfunction ^29^. These diets induce a fat-rich intestinal environment that increases LPS endotoxins, which along with other pathogenic overgrowth, reduces microbial richness and compromises intestinal epithelial integrity that ultimately triggers a systemic inflammatory response ^30^. In contrast, neither MMT nor KGM-MMT hybrid treatments significantly altered Shannon’s index, Chao1 richness, Simpson’s index, or total observed species compared to the HFD group. Although slight increases were noted (∼1.1- to 2.1-fold), these did not reach statistical significance (p > 0.16 for all comparisons). Similarly, Simpson’s index, which reflects species evenness, remained unaffected across treatments (p > 0.08). This suggests that the microbiome restorative effects were primarily attributable to the fermentable properties of KGM as a substrate for the gut microbiota, with MMT exerting minimal independent influence on microbial diversity.

Although spray-dried MMT and KGM-MMT hybrids increased alpha diversity indices (1.1-to 2.1-fold), these changes were not statistically significant (**Fig. 3A**). This aligns with a previous study showing no significant change in the Shannon’s index after five weeks of daily MMT treatment (1 g/kg BW) in HFD-induced obese mice ^11^. As emerging evidence suggests that restoring microbial diversity is protective against metabolic complications in obesity ^27^, the observed KGM-mediated improvements in alpha diversity may confer metabolic benefits for its host.

### 3.4. Modulation of the mice gut micorobiota by KGM based treatments

The microbiota composition analysis further highlights the deleterious effects of a HFD, which enriched harmful pathobionts such as *Streptococcaceae* (**Fig. 4A**). In obese mice, an overgrowth of *Streptococcaceae* was linked to a 2.5-fold increase in fasting insulin levels, a 3-fold increase in serum total cholesterol, and a 3.5-fold increase in liver IL-6 mRNA expression after 8 weeks on a HFD ^31^. In the current study, the dietary inclusion of KGM reduced the mean relative frequency of *Streptococcaceae* by 8.4-log[ folds when compared to the HFD control (p < 0.001) (**Fig. 4B**). Addiotionally, KGM fostered the growth of taxa associated with metabolic benefits, such as *Lactobacillaceae* and *Akkermansiaceae* by 5.7-log[ folds (p = 0.046) and 18-log[ folds (p = 0.0017), respectively. These families, which encompass species like *Lactobacillus rhamnosus* that have demonstrated to enhance intestinal barrier integrity by increasing the expression of tight junction proteins such as zonula occludens-1 (ZO-1) and claudin, are crucial for regulating gut permeability and preventing translocation events across the epithelial barrier ^32^. Another extensively-researched specie from the *Akkermansiaceae* family, *Akkermansia muciniphila,* can activate the nuclear factor kappa-light-chain-enhancer of the activated B cells (NF-κB) pathway in intestinal epithelial cells by releasing metabolites that induce proinflammatory cytokine expression. Simultaneously, it upregulates mucin 2 and tumor necrosis factor alpha-induced protein 3 (TNFAIP3), strengthening the intestinal barrier and reducing inflammation. ^33^.

**Figure 4.**
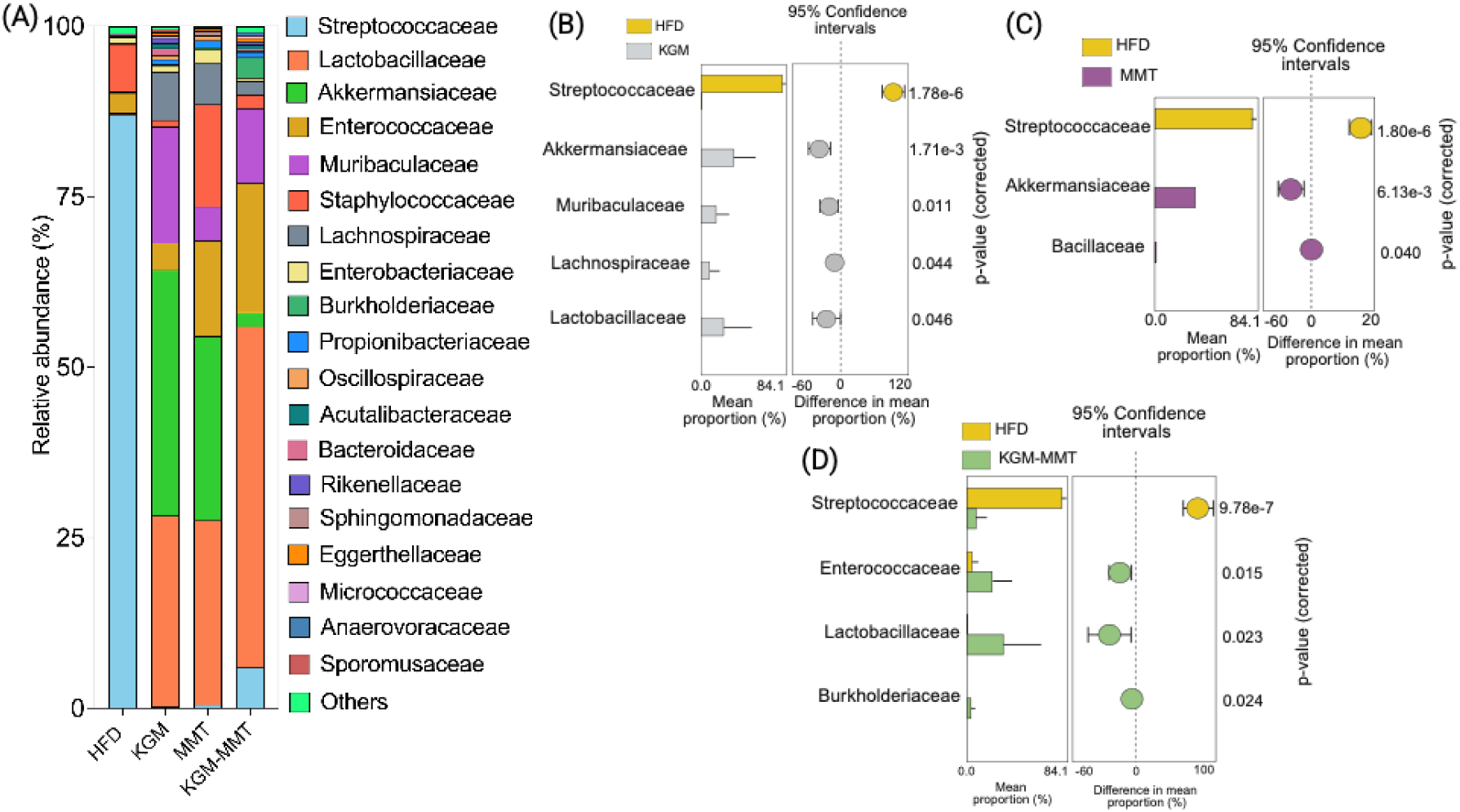
(**A**) Relative abundance of gut microbiota at the family level after the 42-day treatment period across all groups. Differential abundance analysis of family-level taxa comparing the HFD group against (**B**) KGM, (**C**) MMT, and (**D**) KGM-MMT treatments (n = 8 per group). Statistical significance was assessed using Welch’s t-test, with correction for multiple comparisons. Mean relative abundance differences between groups and their associated p-values are shown.

Interestingly, MMT inclusion in the mice diet, also induced similar changes to *Streptococcaceae* (p < 0.001) and *Akkermansiaceae* (p = 0.0061), to those observed with KGM supplementation (**Fig. 4C**). This suggests that MMT may indirectly promote beneficial bacterial growth. One possible mechanism is through adsorption of lipolytic products, thereby lowering luminal lipid levels, which are often linked to dysbiosis ^11^.

In addition the positive microbial shifts in *Lactobacillaceae* (p = 0.023) and *Streptococcaceae* (p < 0.001), KGM-MMT enhanced the relative mean frequency of *Enterococcaceae* by 2.4-log[ fold (p = 0.015) (**Fig. 4D**). This family-level taxa includes species like *Enterococcus cecorum* - linked in HFD-mice studies to improved management of glucose and lipid metabolism disorders ^34^. It is important to recognize that the observed microbial shifts may not be solely due to dietary inclusions. Microbial composition is dynamic and can fluctuate over time, and without longitudinal monitoring, the stability and persistence of these changes remain unclear ^35^. Moreover, while certain family-level taxa are known to include species with metabolic benefits, these families may also harbor species with potentially harmful effects. Future studies should therefore utilize higher-resolution microbial approaches, such as metagenomic sequencing, in order to better postulate the functional consequences of these gut microbiota shifts ^36^.

### 3.5. HFD-induced microbial composition is altered by KGM and MMT treatments

Beta-diversity analysis highlights the differences in microbial composition between treatment groups, providing insight into how gut-active materials affect the gut microbiota of HFD-fed mice. The HFD-fed mice exhibited a significantly distinct microbiome composition compared to all other treatment groups (**Fig. 5A & 5B**). Principal coordinate analysis using Bray-Curtis and Jaccard distance metrics showed that the HFD-fed mice microbiome was significantly different from the KGM (*p* = 0.00006), MMT (*p* = 0.00006), and KGM-MMT (*p* = 0.00018) groups. While the current study demonstrates significant shifts in microbial composition across treatment groups, it is important to note that all groups were subjected to HFD without a normal chow diet control. As such, these microbial alterations cannot solely be attributed to the effects of the HFD itself. Instead, the current results highlight how the different dietary interventions modulate gut microbiota composition within an HFD-fed model. Future studies incorporating a normal diet control group would provide more definitive insights into the extent of HFD-induced microbial shifts. The observed changes in microbial taxa suggest that KGM and MMT supplementation may help mitigate some of the gut microbiota imbalances typically associated with HFD consumption. Previous studies have demonstrated that HFDs induce significant changes in microbial community structure. For instance, an increase in *Firmicutes* and a decrease in *Bacteroidetes* (reported as F/B ratio), are common patterns in microbiome changes driven by HFD, both mice in humans ^37^. Other studies have reported specific enrichments of families such as *Rikenellaceae* and reductions in *Lactobacillaceae* in HFD-fed mice ^38^.

**Figure 5.**
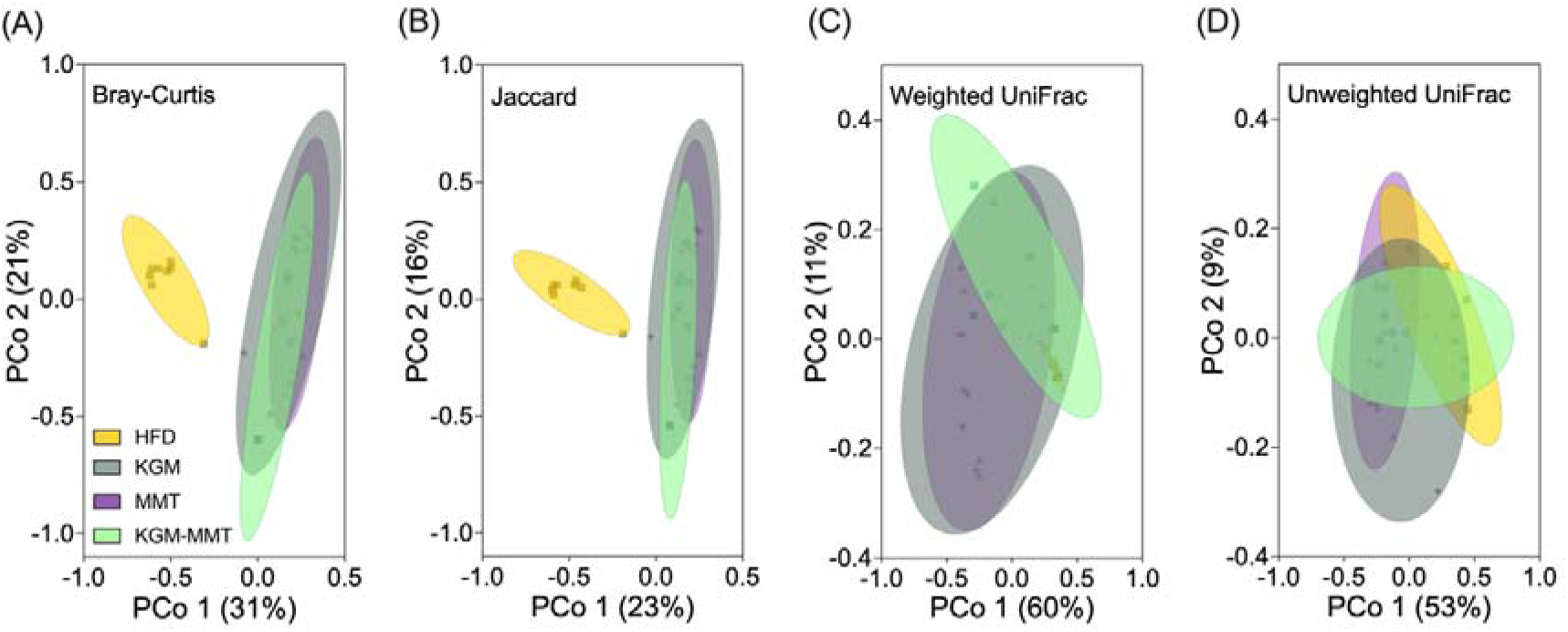
Beta-diversity PCoA plots illustrate the variation in microbial community composition between HFD enriched with KGM, MMT, or KGM-MMT (n = 8 per group). These plots are based on four metrics: **(A)** Bray-Curtis (abundance-based dissimilarity), **(B)** Jaccard (presence/absence-based dissimilarity), **(C)** Weighted UniFrac (phylogenetic distance accounting for relative abundances), and **(D)** Unweighted UniFrac (phylogenetic distance based on presence/absence). PCoA plots with 95% confidence ellipses were used to visualize group clustering and variability. Statistical differences in beta-diversity between groups were assessed using PERMANOVA (permutational multivariate analysis of variance).

The differences between KGM and KGM-MMT were pronouned, with the Bray-Curtis (p = 0.01176) and Jaccard (p = 0.01602) analyses revealing significant diversity differences, suggesting a divergent microbiota emerging between these treatments. Phylogenetic analysis using UniFrac distances, which account for phylogenetic lineages, showed no significant differences between the HFD and treatment groups (**Fig. 5C & 5D**). This indicates that HFD-induced microbiome changes primarily involve shifts within existing microbial lineages rather than the emergence of new ones. For example, taxa such as *Lactobacillaceae* were enriched, and *Streptococcaceae* were depleted with KGM treatment, reflecting a microbial shift within a narrow phylogenetic range. These results contrast other findings that indicate HFD can alter significantly phylum-level changes, however, these changes were most pronouned between 8-12 weeks of HFD-feeding in mice ^39^. Hence, HFD likely alters specific taxa abundances within 6 weeks without broad phylogenetic shifts, while higher-level taxa changes may emerge with prolonged exposure.

### 3.6. Microbial enzymatic diversity enhanced by KGM

Metagenomic predictions derived from 16S rRNA sequences extracted from mice stool samples were conducted using PICRUSt2. These analyses enabled the identification and quantification of enzymatic pathways inferred via the EC database. The results indicate that KGM’s addition to the HFD resulted in a 3.9% to 15% increase in enzyme diversity compared to the HFD group alone (**Fig. 6A**). This is consistent with earlier findings in this study that highlight KGM’s role as a fermentable substrate that promotes gut microbial growth, enhancing microbial enzymatic expression in the process. Similarly, MMT supplementation demonstrated a distinct but less pronounced impact, promoting Shannon’s index by 2.7% (p < 0.05) (**Fig. 6A**). Since MMT can selectively adsorb lipolytic products and influence the microbial environment, it can potentially facilitate the growth of taxa associated with diverse enzymatic functions. However, it is important to note that until now, there have been no published studies which report KGM or MMT functionality using metagenomic predictive tools. Moreover, the hybrid KGM-MMT diet also promoted Shannon’s index by 2.4%, though the increase was not statistically significant (p = 0.0906). This result may reflect a limited study duration, as additive effects between fermentable polysaccharides and MMT could take longer to influence the current HFD-induced mice model of obesity.

**Figure 6.**
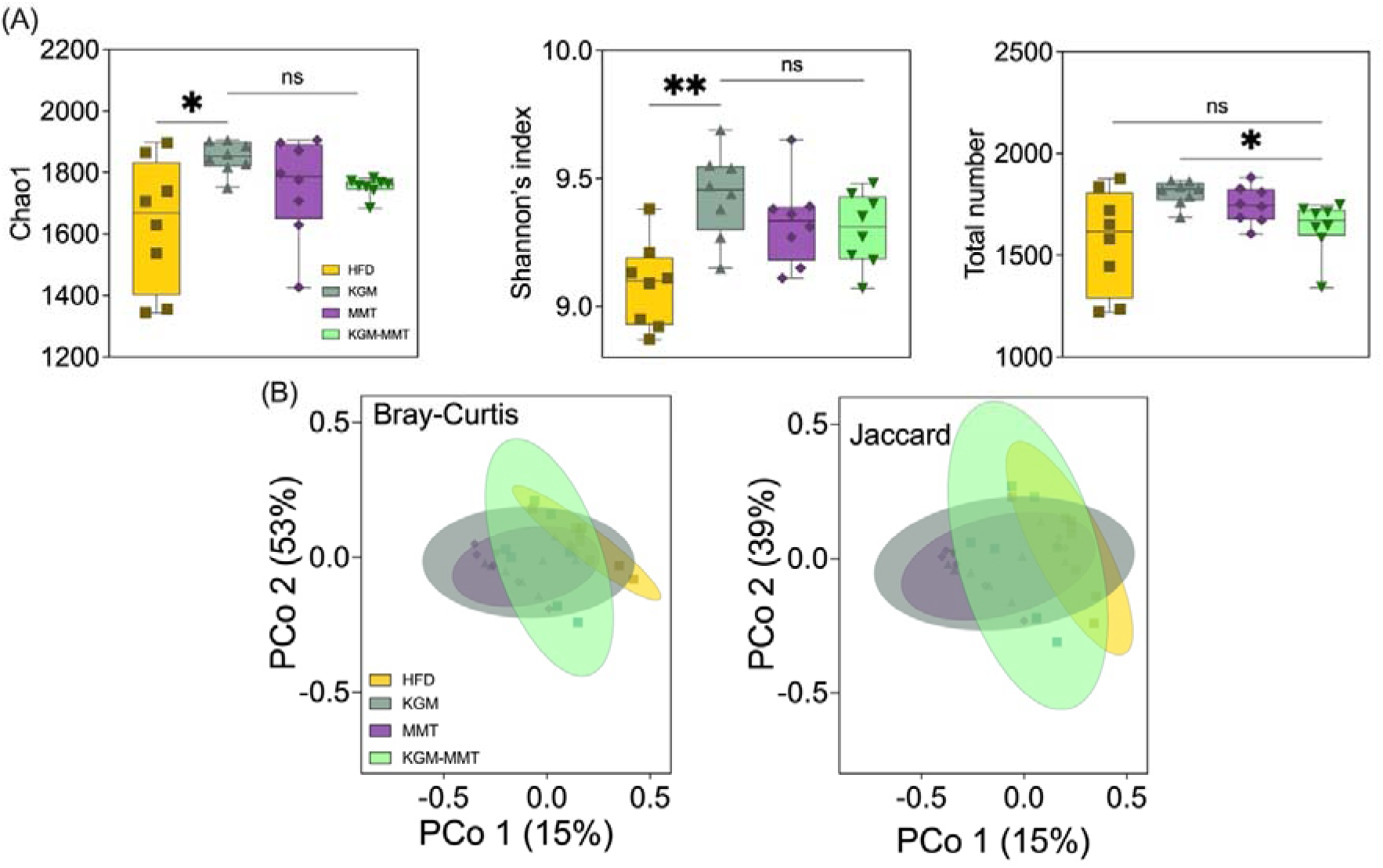
Predicted microbial enzymatic diversity and functional composition across dietary groups. **(A)** Alpha diversity indices, including Shannon’s index and total observed enzyme counts, are presented as box and violin plots displaying min to max values (n = 8 per group), demonstrating significant treatment-induced changes in microbial functional diversity. **(B)** Beta diversity analysis based on Bray-Curtis and Jaccard indices reveals distinct clustering of microbial enzymatic compositions among groups. Statistical differences were determined using the Kruskal–Wallis test with Dunn’s multiple comparisons post hoc (ns > 0.05; *p ≤ 0.05, **p ≤ 0.01, ***p ≤ 0.001.

Beta diversity metrics, Bray-Curtis and Jaccard indices, revealed distinct enzyme compositions among the dietary groups (**Fig. 6B**). Both KGM and MMT diets induced significant shifts in enzymatic composition compared to HFD-fed mice. This supports the study’s findings that KGM and MMT promote a unique gut microbiome composition distinct from that of the HFD-fed mice, which also results in differences in enzymatic profiles from these species. No significant differences in enzyme composition were observed between KGM-MMT and HFD, possibly due to overlapping microbial functions or an insufficient treatment duration.

### 3.7. KGM and MMT enhance enzymes and metabolic pathways regulating intestinal barrier and bile acid function

The analysis of individual enzymes and metabolic pathways revealed significant changes associated with treatments used in this study. As hypothesized, KGM supplementation resulted in a 17-fold increase in the abundance of *mannan endo-1,4-beta-mannosidase* (**Fig. 7A**). This enzyme hydrolyzes beta-1,4 glycosidic bonds in KGM, breaking it into smaller oligosaccharides that gut bacteria can metabolize ^40^. KGM also elevated the relative abundance of *alpha-D-xyloside xylohydrolase*, which cleaves alpha-xylosidic linkages abundant in KGM, by 9.5-fold ^41^. Interestingly, MMT supplementation led to a 6.3-fold increase in the same enzyme, although this was not statistically significant (p = 0.0530). The ability of MMT to influence gut microbial enzymatic activity may be attributed to its role in adsorbing microbial by-products and altering microbial populations, as previously noted in studies examining the effects of natural clays on gut microbiota ^11–13, 15^.

**Figure 7.**
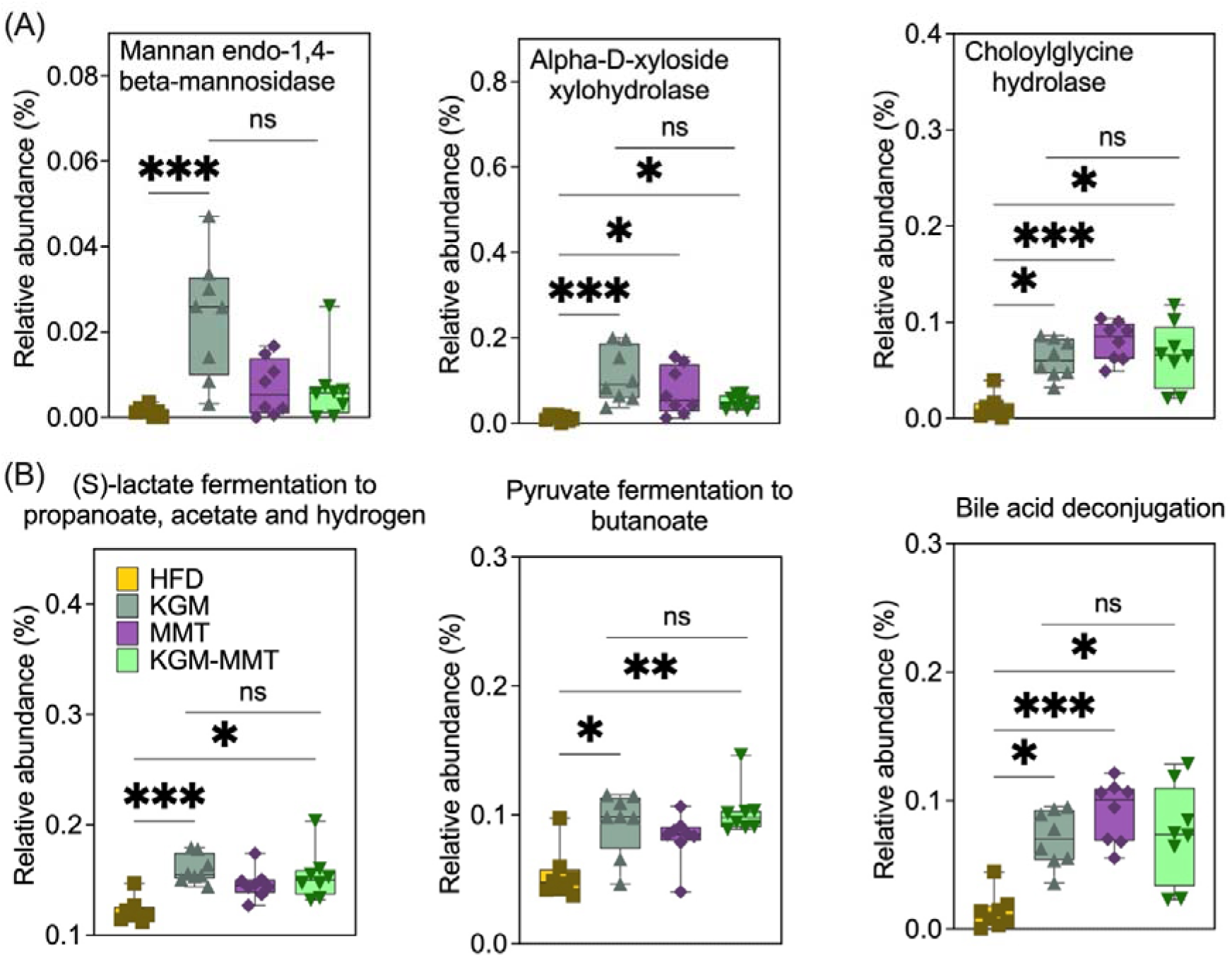
(**A**) Box and violin (minimum to maximum values) plots highlight the relative abundances of key enzymes inferred using the EC database, which show significant increases in *mannan endo-1,4-beta-mannosidase*, *alpha-D-xyloside xylohydrolase* with KGM administration, and *choloylglycine hydrolase* with all treatments (n = 8 per group). (**B**) Metabolic pathways clustered using the MetaCyc database reveal enhancements in SCFA-related fermentation pathways, including (S)-lactate fermentation to propanoate, acetate, and hydrogen, pyruvate fermentation to butanoate, and bile acid deconjugation (n = 8 per group). Statistical differences were determined using the Kruskal–Wallis test with Dunn’s multiple comparisons post hoc (ns > 0.05; *p ≤ 0.05, **p ≤ 0.01, ***p ≤ 0.001.

The relative abundance of *choloylglycine hydrolase* increased by 6.8-fold in mice fed MMT-enriched diets. This enzyme is involved in the hydrolysis of conjugated bile acids, resulting in their deconjugation and thus, restricting their ability to form micelles that facilitate the absorption of dietary lipids ^42^. *Choloylglycine hydrolase* abundance may have been influenced by MMT’s adsorptive properties, which may interfere with bile acid availability. A similar increase was observed in the KGM-MMT hybrid diet (5.6-fold), though this effect was slightly less pronounced, likely due to the reduced proportion of MMT in the hybrid treatment. KGM alone increased *choloylglycine hydrolase* abundance, likely by promoting bile acid–metabolizing bacteria (e.g., *Bifidobacterium spp*. and *Lactobacillus spp.*), with *Lactobacillaceae* significantly enhanced in all treatments in the current study ^43^.

Metabolic pathway analysis revealed notable changes in SCFA production and bile acid metabolism across treatments (**Fig. 7B**). All dietary interventions significantly enhanced the (S)-lactate fermentation to propanoate, acetate, and hydrogen pathway, with increases ranging from 1.2- to 1.3-fold. This pathway is critical for SCFA generation, including acetate, propionate, and butyrate, as well as enhancing gut barrier integrity by promoting the expression of tight junction proteins ^44^. SCFAs can also reduce inflammation by activating G-protein-coupled receptors (GPR41, GPR43) and inhibiting histone deacetylases ^45^. These actions help modulate immune responses and reduce chronic low-grade inflammation often seen in obesity ^46^. KGM administration resulted in the greatest increase in this pathway, followed by the KGM-MMT hybrid. This is consistent with the role of KGM as a fermentable polysaccharide, promoting the microbial production of SCFAs. Similarly, pyruvate fermentation to butanoate was significantly enhanced across all treatments, with increases ranging from 1.5- to 1.9-fold. The KGM-MMT hybrid produced the greatest enhancement, potentially due to the additive effects of MMT and KGM on microbial activity. Butyrate, a key product of this pathway, is a crucial energy source for colonocytes, fulfilling 70-80% of their energy requirements through oxidation, which also helps maintain an anaerobic environment favorable for beneficial microbiota ^47^. Moreover, butyrate administration can decrease pro-inflammatory cytokines (e.g., IL-1β, IL-6, IL-8, TNF-α), whilst simultaneously promoting anti-inflammatory cytokines (e.g., IL-10, TGF-β) ^48^.

Bile acid metabolism was also significantly influenced, with bile acid deconjugation enhanced by 5.1-, 5.4-, and 6.7-fold for KGM, KGM-MMT, and MMT treatments, respectively. The potent effects of MMT-based treatments may be due to their role in bile acid restriction, reducing dietary fat availability in the process ^11–13, 15, 16^. KGM also contributed to increased bile acid deconjugation, likely through its fermentation by the gut microbiota, which promotes microbial enzymes capable of bile acid metabolism ^43^. The less pronounced increase observed with the KGM-MMT hybrid can be due to the lower proportion of the clay in the treatment, highlighting the dose-dependent effects of MMT on bile acid modulation. Importantly, the

current enzymatic and metabolic predictions suggest MMT’s adsorptive properties selectively favor microbial enzymes involved in bile acid deconjugation.

### 3.8. KGM-MMT reduces HFD-induced weight gain

After 42 days of dietary intervention, all treatment groups exhibited reduced body weight gain compared to HFD-fed controls (**Fig. 8A**). Area under the curve (AUC) analysis of body weight change confirmed this pattern, with all treatments significantly reducing cumulative body weight gain compared to HFD (p < 0.0001 for all comparisons). Notably, the KGM-

**Figure 8.**
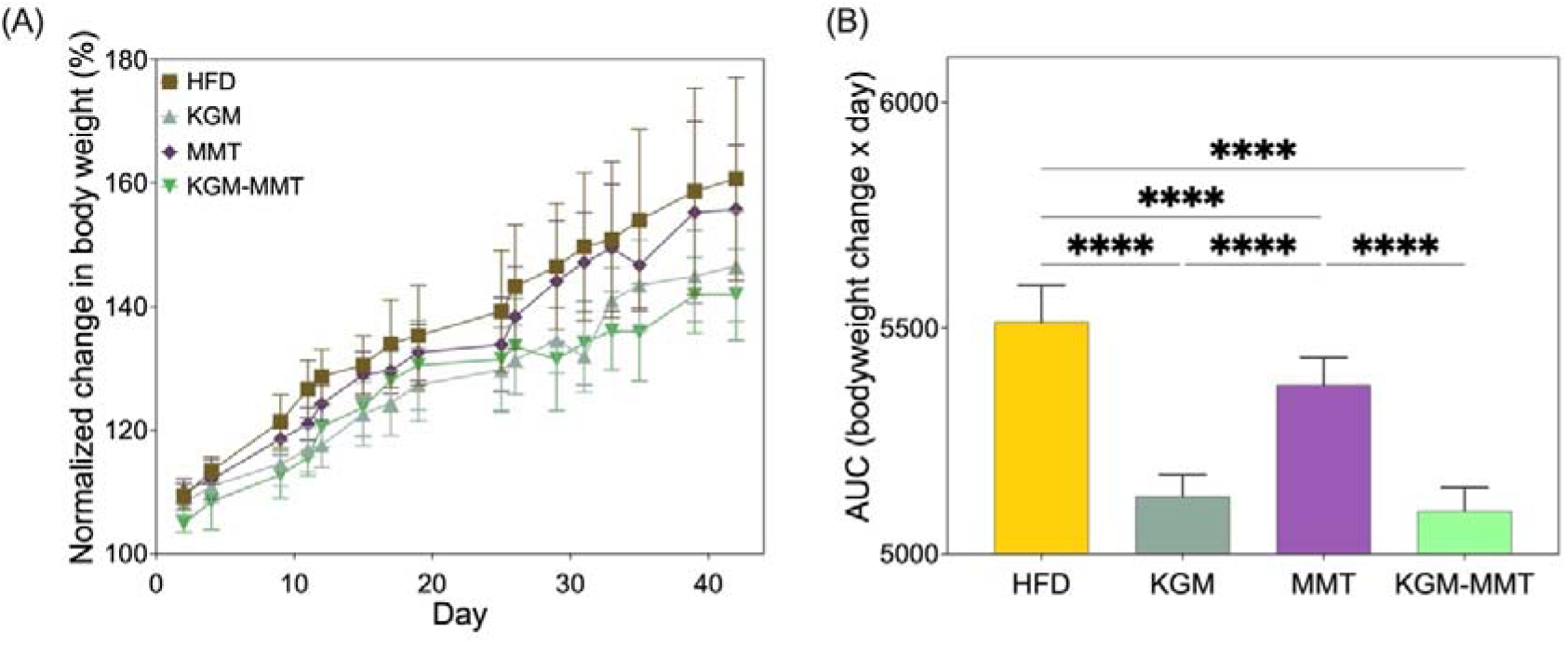
(**A**) Normalized bodyweight changes over 42 days show reduced weight gain with all treatments compared to HFD (n = 8 per group). (**B**) Area under the curve (AUC) analysis confirms the greatest reduction in weight gain with the KGM-MMT hybrid, followed by KGM and MMT treatments. Statistical differences were determined using one-way ANOVA with Tukey’s comparisons post hoc (ns > 0.05; *p ≤ 0.05, **p ≤ 0.01, ***p ≤ 0.001, ****p ≤ 0.0001).

MMT hybrid also outperformed MMT (p < 0.0001), though it did not significantly differ from KGM alone (p = 0.6116) (**Fig. 8B**). Weight gain profiles across the treatment phase remained similar across groups until approximately day 25 (**Fig. 8A**). After this point, mice receiving KGM and KGM-MMT displayed a distinct plateau in body weight gain, whereas those in the HFD and MMT groups continued to gain weight steadily. These results align with prior findings that weight divergence between standard chow and HFD-fed rodents becomes evident after 3–4 weeks of dietary intervention ^49^. KGM’s anti-obesity effects have been widely reported and are attributed to mechanisms such as delayed gastric emptying, increased gut viscosity, and stimulation of satiety hormones including GLP-1 and PYY ^24^. Recent findings further show that KGM activates thermogenesis in inguinal white adipose tissue via β3-adrenergic receptor (ADRB3) signaling, increasing UCP1 expression and promoting fat oxidation ^50^.

In contrast, MMT primarily acts by adsorbing dietary lipids, bile acids, and lipolytic products in the gut, thereby reducing fat absorption and its bioavailability ^11, 13, 14^. While this mechanism contributed to a modest reduction in weight gain, MMT was notably less effective than KGM or the KGM-MMT hybrid. Among all groups, the KGM-MMT hybrid demonstrated the greatest reduction in AUC (5094[±[52.95), corresponding to a 7.6% decrease compared to HFD (p < 0.0001) and a 5.2% decrease compared to MMT (p < 0.0001). Although the reduction was not statistically different from KGM alone, the hybrid still delivered the lowest cumulative weight gain. This suggests a potential additive effect, likely due to the combination of KGM’s satiety-enhancing and thermogenic properties with MMT’s capacity to reduce fat absorption. This multi-mechnastic action for hybrids are supported by similar findings in other fiber-based hybrid systems. For instance, bacterial cellulose–KGM and KGM– dihydromyricetin complexes have demonstrated enhanced metabolic effects, including greater reductions in body weight, insulin resistance, and oxidative stress, than either component alone ^3, 51^. However, an important limitation of the present study is the absence of a standard chow diet control group. Without this reference, it is not possible to determine whether the dietary interventions normalized weight gain toward metabolically baseline levels, or only attenuated the degree of HFD-induced gain. Future studies should include chow-fed controls to fully evaluate the potential of these interventions to restore metabolic health.

### 3.9. KGM -MMT reduces inflammation and blood glucose whilst improving body composition

All treatment groups showed marked reductions in serum IL-6 concentrations relative to HFD-fed controls (**Fig. 9A**). IL-6 levels were reduced by approximately 95% in the MMT group (p = 0.0030) and 97% in the KGM-MMT group (p = 0.0002). Although the KGM group also demonstrated a substantial 93% reduction, this did not reach statistical significance (p = 0.0555). These reductions are consistent with previous studies showing that KGM and MMT downregulate inflammatory signaling in HFD-induced obesity ^3, 11, 12^. KGM, particularly in fiber hybrid forms, has been shown to lower hepatic IL-6 and TNF-α by 39% and 53%, respectively ^3^, while MMT reduces hepatic IL-6, TNF-α, and COX-2 expression, restoring inflammatory markers toward healthy levels ^11, 12^. Serum IFN-γ levels, however, did not differ significantly across groups in the current study (**Fig. 9B**). While MMT and KGM-MMT demonstrated reductions (∼27–29% below HFD), these changes were not statistically significant (p > 0.05 for all comparisons). This suggests a selective anti-inflammatory effect, with IL-6 more sensitive to dietary modulation than IFN-γ in this model. Future studies should therefore assess a broader inflammatory profile, including cytokines such as TNF-α, IL-10, and MCP-1, to better characterize systemic immune responses and their relationship with the current treatments.

**Figure 9.**
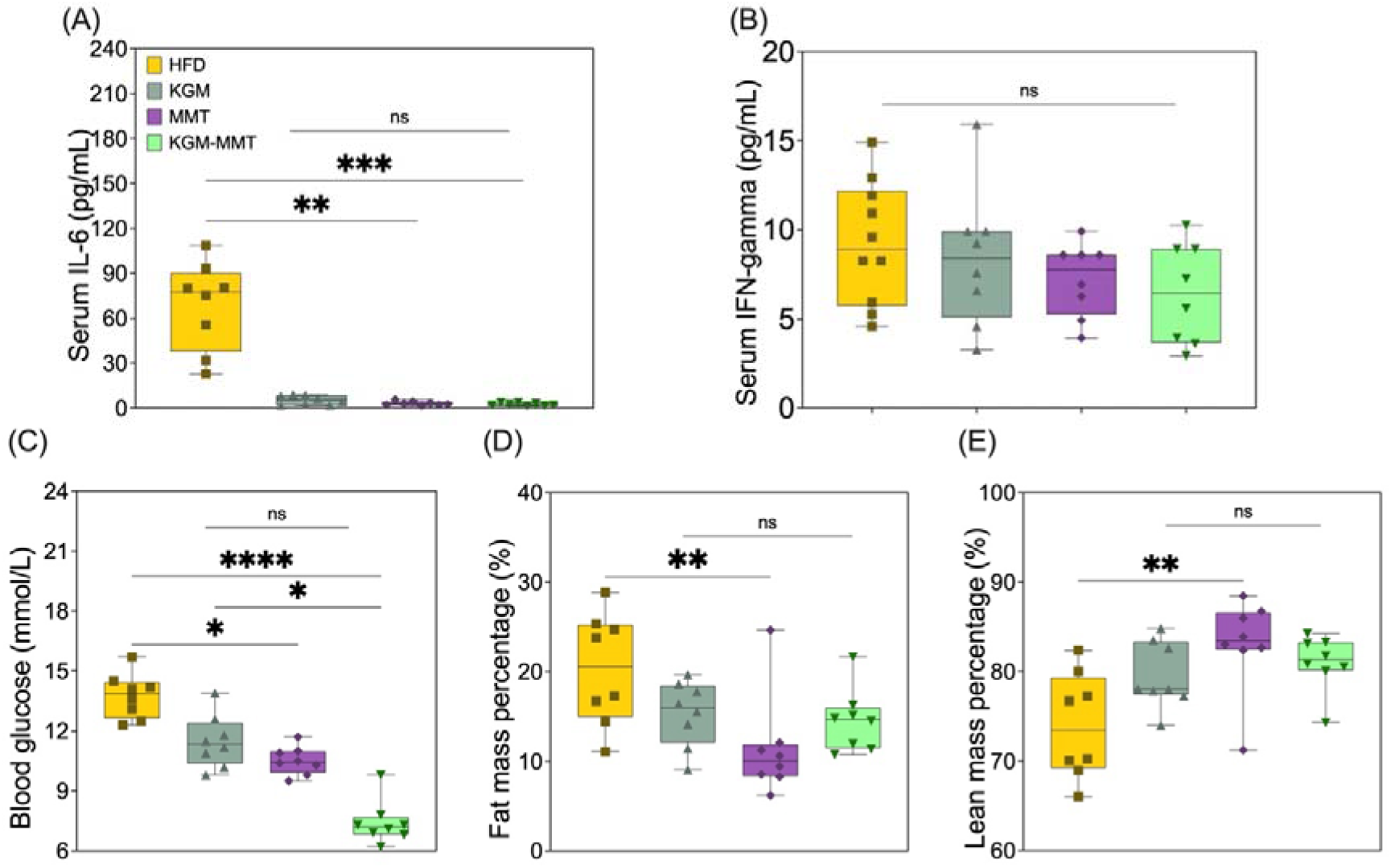
Effects of dietary treatments on systemic inflammation, blood glucose, and body composition in HFD-fed mice after 42 days. (**A**) Serum IL-6 concentrations were significantly reduced by 95% with MMT and by 97% with KGM-MMT. KGM reduced IL-6 by 93% but was not significant (p = 0.0555). (**B**) Serum IFN-γ levels were not significantly different across groups. (**C**) Blood glucose levels decreased by 16% (KGM), 24% (MMT, p = 0.0271), and 46% (KGM-MMT, **p < 0.0001). (**D**) Fat mass percentage was significantly reduced by 44% with MMT (p = 0.0076), while the 29% reduction in the KGM-MMT group was not significant. (**E**) Lean mass increased by 12% with MMT (p = 0.0032) and by 10% with KGM-MMT (ns).

Blood glucose levels were significantly reduced in the MMT group by 24% (p = 0.0271) and in the KGM-MMT group by 46% (p < 0.0001) compared to HFD-fed controls (Fig. 9C). The KGM group resulted in a 16% reduction that did not reach statistical significance (p = 0.3831). Moreover, glucose levels in the KGM-MMT group were significantly lower than those in the KGM group (p = 0.0136), suggesting a potential additive effect. These effects align with established mechanisms, whereby KGM delays gastric emptying and enhances insulin signaling via upregulation of glycolytic enzymes and glycogen synthesis ^4, 52^, while MMT reduces nutrient absorption by binding dietary lipids and bile acids ^11, 12^. In terms of body composition (**Fig. 9D & 9E**), MMT significantly reduced fat mass by 44% (p = 0.0076), while KGM-MMT reduced it by 29% (p = 0.5908). Both groups showed increases in lean mass, with MMT producing a 12% gain (p = 0.0032) and KGM-MMT a 10% gain (p = 0.0988). These data suggest that while KGM-MMT may not significantly improve body composition beyond MMT, it preserves these benefits while enhancing glycemic and inflammatory outcomes. Collectively, these results suggest that the KGM-MMT hybrid combines complementary mechanisms—KGM’s effects on satiety and glycemic control, and MMT’s lipid-binding and adiposity-limiting functions—to produce broad metabolic benefits.

Data shown as box and violin plots (min–max). n = 8 per group. Statistical differences were determined using the Kruskal–Wallis test with Dunn’s multiple comparisons post hoc (ns > 0.05; *p ≤ 0.05, **p ≤ 0.01, ***p ≤ 0.001.

### 3.10. KGM-MMT hybrids reduce food intake and enhance satiety

During the final week of the 42-day treatment period, mice were housed in metabolic cages to monitor feeding behavior and metabolic parameters. All treatment groups demonstrated reductions in total daily food intake compared to HFD-fed controls (**Fig. 10A**). Mean daily intake was reduced by 65% with KGM (p = 0.0029), 63% with MMT (p = 0.0047), and 51% with KGM-MMT (p = 0.0681). Although the KGM-MMT group consumed slightly more than the individual components, the difference was not statistically significant.

**Figure 10.**
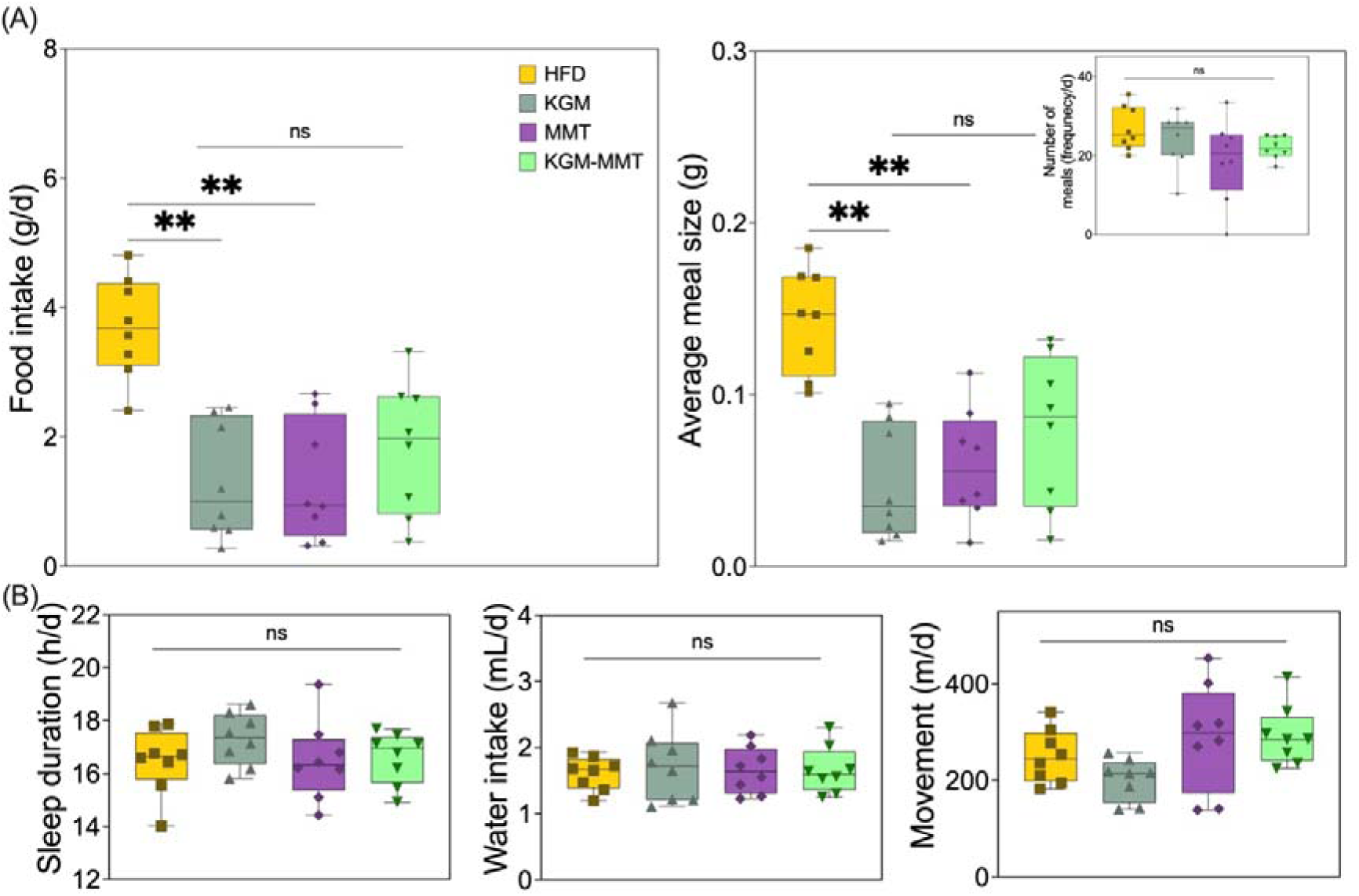
(**A**) Box and violin (min–max) plots show that all treatments reduced food intake by 51–65% and meal size by 45–66%, without altering meal frequency (inset), suggesting enhanced satiety (n = 8 per group). (**B**) Sleep duration, water intake, and movement were unaffected, supporting the hypothesis that changes in feeding behavior are not stress-induced but due to physiological effects of the interventions. Statistical differences were determined using the Kruskal–Wallis test with Dunn’s multiple comparisons post hoc (ns > 0.05; *p ≤ 0.05, **p ≤ 0.01, ***p ≤ 0.001.

Average meal size followed a similar pattern. Reductions of 66% (KGM) (p = 0.0012) and 58% (MMT) (p = 0.0087) were observed, while the KGM-MMT group showed a 45% decrease that was not statistically significant (p = 0.1409). Notably, meal frequency remained unchanged across all groups (data not shown), suggesting that reduced intake was due to enhanced satiety rather than behavioral suppression or altered feeding patterns. Previous studies have shown that KGM promotes satiety via increased gut viscosity, delayed gastric emptying, and elevated levels of satiety-related hormones such as Glucagon-like peptide-1 (GLP-1) and Peptide YY (PYY) ^24, 53^. MMT may act via complementary or similar pathways. Although MMT’s impact on gastric viscosity remains unexplored, it has been demonstrated to modulate the gut microbiota, increasing taxa such as *Akkermansia* that influence tryptophan and arginine metabolism, both linked to appetite regulation ^54^. These effects could theoretically enhance satiety signaling, although this was not directly measured in the current study. Importantly, no significant differences were observed in behavioral stress indicators including sleep duration, water intake, or movement activity (**Fig. 10B**, all p > 0.05). This suggests that reduced food and meal size were unlikely due to stress or altered activity and more likely reflect treatment-induced physiological effects. It must be acknowledged that the KGM-MMT hybrid did not significantly outperform KGM or MMT alone in reducing food intake or meal size. Moreover, while reduced intake and meal size support a satiety-enhancing role, the mechanistic basis remains unknown. Future studies should therefore, directly investigate the underlying mechanisms through gastric emptying assays, as well as systemic hormonal markers (e.g., GLP-1, PYY).

## Conclusions

This study highlights the therapeutic potential of a spray-dried KGM-MMT hybrid as a novel dietary intervention for improving metabolic health in the context of HFD-induced obesity. By integrating KGM’s fermentable, satiety-promoting properties with MMT’s lipid-adsorptive and anti-inflammatory effects, the hybrid produced an overall additive benefit across multiple metabolic markers, including reductions in serum IL-6 and blood glucose, improvements in lean mass, and suppression of food intake and meal size. Although KGM-MMT did not consistently outperform KGM or MMT alone all measurments, overall, it retained the therapeutic benefits of both, supporting its utility as a single-dosage form used to attenuate diet-induced metabolic effects. As such, the use of spray-drying to fabricate KGM-MMT microparticles introduces a scalable application for delivering multifunctional metabolic effects via gut-targeted mechanisms. Unlike standalone biomaterials, KGM-MMT hybrid offers the potential to modulate inflammation, appetite, and energy metabolism concurrently, using components that are food-grade. While further studies are warranted to explore long-term efficacy, dose response, and mechanistic pathways, the current findings elucidate novel insights for the development of engineered fiber–clay formulations as the next-generation of adjuvant strategies for metabolic disease management.

## Conflicts of interest

The authors declare no conflict of interest.

## Acknowledgments

P.J. acknowledges the Channel 7 Children’s Research Foundation (21-16816523) and the Hospital Research Foundation (THRF) group (2022-CF-EMCR-004-25314) for their generous support of this project. H.R.W. would like to thank the Hospital Research Foundation Group and NHMRC for their ongoing support in the form of research fellowships.

## Data Availability Statement

The data for all findings of the current study is available at request from the corresponding author.

## Notes

### Competing Interest Statement

The authors have declared no competing interest.

